# Immediate effects of psychosocial stress on attention depend on subjective experience and not directly on stress-related physiological changes

**DOI:** 10.1101/223909

**Authors:** I. Palacios-García, M. Villena-Gonzalez, G. Campos-Arteaga, C. Artigas-Vergara, K. Jaramillo, V. López, E. Rodríguez, J. R. Silva

## Abstract

Acute psychosocial stress is associated with physiological, subjective and cognitive changes. In particular, attention, which is considered one of the main processes driving cognition, has been related to different stress outcomes, such as anxiety, cortisol levels and autonomic responses, individually. Nonetheless, their specific contributions to and association with attention is still not fully understood. To study this association, 42 male participants were asked to perform an attentional task just before and immediately after being exposed to either an experimental treatment designed to induce psychosocial stress using the Trier Social Stress Test (TSST) or a matching stress-free control condition. The salivary cortisol concentration, heart rate, and self-reported anxiety were measured to assess the physiological response to stress and the subjective experience during the protocol. As expected, psychosocial stress induced increases in heart rate, salivary cortisol levels and anxiety. The behavioral analysis revealed that members of the control group performed better on the attentional task after the protocol, while members of the TSST group showed no changes. Moreover, after dividing the stress group into sub-groups of participants with high and low anxiety, we observed that participants in the high-anxiety group not only failed to perform better but also performed worse. Finally, after testing several single-level mediation models, we found that anxiety is sufficient to explain the changes in attention and that it mediates the effects between heart rate and cortisol levels on attention. Our results suggest that the immediate effects of acute psychosocial stress on attention are highly dependent on the participant’s subjective experience, which, in turn, is affected and can mediate stress-related physiological changes.

## Introduction

Many daily life experiences are characterized by unpredictable and uncontrollable situations in which the individual anticipates psychological consequences, which are generally produced by social factors. Those situations are responsible for psychosocial stress (Kudielka, 2007). The psychosocial stress response includes a physiological component characterized by the activation of the hypothalamic pituitary adrenal axis (HPAA) and the autonomic nervous system (ANS), resulting in increased cortisol release and elevated heart rate, respectively. Changes at the physiological level have been shown to be closely related to cognitive performance, especially working memory (Chamberlain, Muller, Blackwell, Robbins, & Sahakian, 2006), memory retrieval (Kuhlmann, Kirschbaum, & Wolf, 2005) and selective attention (Cornelisse, Joels, & Smeets, 2011).

In addition to physiological responses, social stress is associated with a psychological component, such as an emotional experience, the perception of stress or an increase in the self-reported state of anxiety (Hellhammer & Schubert, 2012). In particular, a state of anxiety can be conceptualized as “the state in which an individual is unable to instigate a clear pattern of behavior to remove or alter the event/object/interpretation that is threatening an existing goal’’ (Power & Dalgleish, 2015). Individuals who are not anxious are generally able to select the information required to perform a specific task according to their goals; this process is referred to as attentional control (Gazzaniga, 2004; Petersen & Posner, 2012; Serences et al., 2005). Nevertheless, higher levels of anxiety may affect this process. Concordantly, Eysenck, Derakshan, Santos, and Calvo (2007) developed attentional control theory and discussed how anxiety may affect attentional control and cognitive performance. Several researchers have addressed this issue; for instance, Harris and Cumming (2003) demonstrated that individuals with high levels of anxiety did not perform a task involving prospective memory, i.e., one that involves recalling a planned intention at some future point in time. Furthermore, it has been suggested that an elevated level of anxiety impairs task switching, resulting in more errors and increasing the task completion time (Goodwin & Sher, 1992).

As previously shown, both physiological and psychological responses are implicated with regard to cognitive/behavioral changes. However, during a psychosocially stressful situation, both responses interact and lead to a mixture of outcomes that can vary depending on the type, timing and severity of the stressor and the task (Allen, Kennedy, Cryan, Dinan, & Clarke, 2014). One of the most common experimental settings for inducing stress is the Trier Social Stress Test (TSST) (C. Kirschbaum, Pirke, & Hellhammer, 1993). Studies using the TSST have revealed some of the impairments that social stress may produce at different cognitive levels, including working memory (Schoofs, Preuss, & Wolf, 2008), attention (Olver, Pinney, Maruff, & Norman, 2014), multitasking (Plessow, Schade, Kirschbaum, & Fischer, 2012) and second- and higher-order processes, such as metacognition (Reyes, Silva, Jaramillo, Rehbein, & Sackur, 2015).

Although the majority of studies using the TSST have reported physiological and psychological responses, the relationship between both responses in the context of a cognitive task is not yet fully understood. Interestingly, a recent study (Ali, Nitschke, Cooperman, & Pruessner, 2017) proposed the use of a pharmacological procedure to study the effects of the TSST on participants’ psychological responses in the absence of physiological responses. The results showed that suppressing the physiological response did not impact the emotional response to the stressor, giving rise to the question of whether the physiological stress response in fact contributes to the emotional experience. The latter study, however, did not include a cognitive task. Similar caveats can be found in other studies of the relationship between the psychological perception of stress and cognition (task-switching), which do not consider the physiological component (Liston, McEwen, & Casey, 2009), and studies that show strong interdependency between physiological response (cortisol reactivity) and cognitive performance (selective attention), but do not assess the role of psychological stress response (Roelofs, Bakvis, Hermans, van Pelt, & van Honk, 2007). There is strong evidence linking both the physiological and psychological responses related to social stress with cognition. However, the relationship between these variables with regard to a specific behavioral/cognitive outcome is still not understood. Moreover, in the study of attention, there is evidence that anxiety is a relevant factor; however, few studies have connected anxiety with stress-dependent cognitive changes.

We aimed to study—in the same experimental context—the relationship between physiological response, psychological experience and the cognitive performance associated with psychosocial stress. Specifically, we studied the relationship between cortisol reactivity, heart rate, self-reported state of anxiety and performance in an attentional task in individuals exposed to a stress protocol (TSST) or a control protocol.

We hypothesized the following: (1) that psychosocial stress impairs an individual’s performance in an attentional task; (2) that the physiological response—measured using the salivary cortisol concentration and the heart rate—is positively correlated with the self-reported state of anxiety; and (3) that, in accordance with attentional control theory, the psychosocial-stress-dependent increase in the state of anxiety—and not in the physiological response—mediates the participant’s performance impairment in the attentional task.

## Methods

### Participants

Forty-nine male participants were recruited and assigned to the stress (n = 24) or control (n = 25) group. Seven of these participants (4 controls) were excluded because they failed to follow instructions during the experiment or because of data acquisition problems. The outcomes of the 42 healthy non-medicated volunteers (mean age, 25 ± 3.8 years) were recorded between 12:00 am and 2:30 pm; the participants were instructed to avoid smoking cigarettes, drinking coffee or tea and eating food, including bubble gum, 2 hours before the experiment. All the participants provided written informed consent before the study in accordance with the guidelines of the Bioethics Committee of the Faculty of Medicine at Pontificia Universidad Católica de Chile, which approved the research protocol.

### Procedures

The participants underwent electroencephalogram (EEG), electrocardiogram (EKG) and electrooculogram (EOG) electrode placement (the EEG and EOG data are not shown). The experiment commenced with an initial attentional task (pre-condition) flanked by 90 seconds of resting state recording. Subsequently, the TSST or the control protocol was conducted, and the flanked attentional task (post-condition) was repeated (Figure 1). In addition to attentional performance, heart response and salivary cortisol levels were monitored to assess the activation of the ANS and the HPAA, respectively. The EKG was registered as an external electrode in the EEG recording setup using a BioSemi ActiveTwo® system throughout the experiment. Salivary samples for cortisol measurement were obtained at 5 different times (Section Method/Physiological measurements). Finally, the participants were asked to report their levels of anxiety just before and after the TSST or control protocol. The number of interruptions (cortisol sampling and the anxiety scale) during the procedure was minimized to maintain the participants’ involvement and the natural physiological and subjective states induced by the protocol.

**Figure 1.**
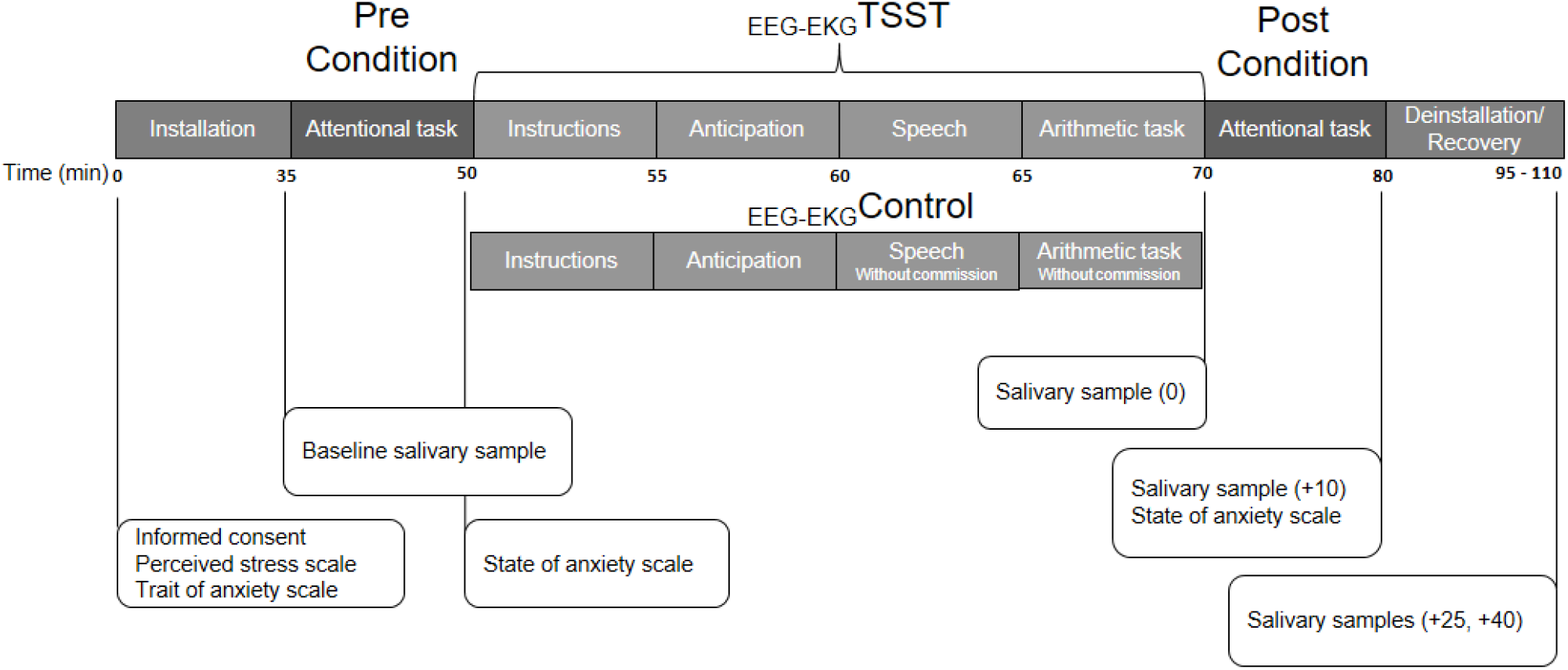
General experimental protocol showing all the data collected over time. Numbers indicate time in minutes.

### Stress induction and control protocol

Psychosocial stress was induced using an EEG-adapted version of the Trier Social Stress Test (C. Kirschbaum et al., 1993). The protocol consisted of a simulated interview in which the participants must provide personal information to apply to a fictional job (5 minutes) in front of three people acting as referees (who appeared serious and expressionless) and a video camera. This was followed by an arithmetic task (5 minutes) consisting of subtracting 13 from 1000 repeatedly until 0; each time the participant made a mistake, he was told by one of the referees to start over. The protocol follows the guidelines proposed by C. Kirschbaum et al. (1993) except that the referees were the ones who entered the room while the participant prepared for the evaluation inside the room.

The control protocol included the same procedures except that the subjects performed in front of the experimenter (who appeared to be in a good mood and who maintained a friendly attitude) instead of people acting as referees. The same physical and mental effort was induced but the psychosocial stress component was removed. After the protocol, the participants were informed that no judgments were made about their presentation and that the camera was turned off.

### Attentional task

An adaptation of a task-switching paradigm by Liston, Matalon, Hare, Davidson, and Casey (2006) was used as the attentional task. Two circles, each subtending 4.6° of the visual space and equidistant of the monitor’s center, were presented for 700 ms. Each circle was red or green and moved upward or downward. Between them, there was a letter appeared: “M” for movement or “C” for color. The subject was instructed to choose the green circle when the letter was “C” and the upward circle when the letter was “M.” Each trial began with a central white fixation cross of variable duration (600–1000 ms). The complete trial involved the central fixation followed by 700 ms of the colored and mobile circles. Accuracy and reaction times were recorded on a trial-by-trial basis using PsychoPy (Peirce, 2008). The participants were trained using three blocks of 12 trials requiring color, movement and color/movement discrimination. The experiment involved four blocks of 64 trials separated with 1-minute rests between blocks.

We modified the task-switching paradigm (Liston et al., 2006) by approximately doubling the task speed. This change was made to ensure that the mean accuracy was 70%, which required the task to be substantially more difficult than previously reported.

### Physiological measurements

EKG activity was monitored during the sessions using 2 external electrodes (BioSemi ActiveTwo®) and placing 2 fingers under the left collarbone and over the left hip to observe the R peaks during a specific time period and measure the heart rate during this period. Seven different 90-second periods were used to calculate the heart rate during the four rest periods and at the beginning of different control/TSST tasks (anticipation, speech and math) (Figure 1). We decided to consider only the first 90 seconds of the control/TSST tasks to equate the variability produced by comparing periods with different lengths. The heart rate was obtained and calculated using custom MATLAB scripts and Kubios (Tarvainen, Niskanen, Lipponen, Ranta-Aho, & Karjalainen, 2014).

In addition to the heart response, 1 mL of saliva was collected in a salivary tube at five different time points (Figure 1): just before the control or TSST treatment (baseline) and 0 (+ 0), 10 (+ 10), 25 (+ 25) and 40 minutes (+ 40) after the beginning of the control or TSST treatment. Immediately after collection, the samples were preserved at −20°C until they were analyzed. The saliva samples were sent to the Molecular Biology Laboratory of the Universidad de La Frontera for quantitative measurements of the cortisol concentration. The salivary concentrations of cortisol were obtained using an enzyme immunoassay commercial kit following the manufacturer’s instructions (DRG Salivary Control ELISA Kit, DRG Instruments GmbH, Germany) (Reyes et al., 2015).

### Psychological response

The perceived stress scale (Cohen, Kamarck, & Mermelstein, 1983) and the trait anxiety scale (Spielberger & P. R., 1983) were used before each procedure to assess the daily and baseline subjective stress state and trait anxiety, respectively. Regarding the psychological experience of our experimental design, the participants were asked to report their level of anxiety using the scale (Spielberger & P. R., 1983) just after and before the control or TSST protocol (Figure 1). The psychological response associated with our experiment focused exclusively on the acquisition of stress-dependent self-reported anxiety.

### Single-level model

We studied the relationship between salivary cortisol, heart rate, self-reported level of anxiety and attentional performance by testing different single-level models. The model implementation involves initial (X), outcome (Y) and mediator (M) variables. In other words, there is an initial independent variable X that better explains the dependent outcome variable Y when the mediator variable M is present (Baron & Kenny, 1986). The mediated effect was studied by estimating the regression equation that predicts the outcome Y from the initial variable X,

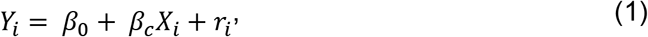

and by estimating the regression equation that predicts the outcome Y from the initial variable X and the mediated variable M,

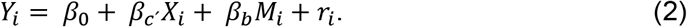

The difference of the coefficients associated with the initial variable X (c – c’) reflects the mediation effect of the variable M on the predicted effect of the variable X on the outcome Y.

Additionally, the mediation effect can be estimated by estimating the regression equation, which predicts the mediator M from the initial variable X, by multiplying the regression coefficient of the mediator M explaining the outcome Y (*b*) from equation (2) by the regression coefficient of the initial variable X explaining the mediator M (*a*) from equation (3), we obtain a second estimate of the mediation effects (*a* * *b*), (Krull & MacKinnon, 2001).

Finally, the mediation estimation can be calculated using

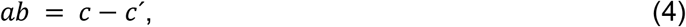

where *a* is the coefficient of the initial variable X predicting the mediator M, *b* is the coefficient of the mediator M predicting the outcome Y, *c* is the coefficient of the initial variable X predicting the outcome Y and c’ is the coefficient of the initial variable X predicting the outcome Y in the presence of the mediator M^1^ (Baron & Kenny, 1986).

We used the single-level functions implemented on MATLAB’s Multilevel Mediation/Moderation Toolbox (M3) v.0.9 (T.D.W. and M.A.L). The toolbox calculates the first-level path coefficients (PC) using standard ordinary least-squares multiple regression and determines p values using the bootstrapping approximation principle (for additional details, see the supplementary method of Wager et al. (2009)).

This study was designed to assess the relationship between stress levels—measured using salivary cortisol levels, heart rates and levels of anxiety—and the changes in attentional performance in the control and experimental protocols. Because there is no clear and defined relationship between these variables and attention, six different models using attentional performance as outcome Y and choosing the initial variable X and the mediator M from among the salivary cortisol level, heart rate and level of anxiety were tested. Therefore, each variable was considered the initial variable X while the other two variables comprised the mediator. For each model, the a, b, c, c’ and ab coefficients were obtained. The significance of each coefficient was tested using bootstrapping (n = 10000).

For the analysis, the following values were calculated for the 42 participants and each model: a unique value of each variable, the change in salivary cortisol (delta; the level after each protocol minus the baseline level), the levels of anxiety and attentional performance, and the area under the curve for the heart rate response.

### Statistical analysis

The differences between the groups in terms of daily perceived stress and trait anxiety were evaluated using a two-tailed unpaired t-test. The effects of psychosocial stress (the TSST) on attentional performance, physiology and the level of anxiety were evaluated using a two-way repeated-measures analysis of variance (ANOVA) together with the Bonferroni post-test.

The relationship between the changes in attentional performance and anxiety was evaluated using Pearson’s correlation. All the analyses were performed using GraphPad Prism (GraphPad Software, San Diego CA, USA). The data in the graphs are presented as mean ± SEM.

## Results

### Physiological and psychological stress responses

We measured the heart rate, salivary cortisol level and self-reported level of anxiety to corroborate the success of our experimental and control designs. The participants reported increased levels of anxiety only after the stress protocol (Figure 2A; Group × Time interaction; F(1,41) = 42.03; p<0.001. Bonferroni post hoc test; Pre-Treatment, *t* = 2.122, p>0.05. Post-Treatment, *t* = 8.493, p<0.001), whereas after the control protocol, the level of anxiety was nearly unaffected. Additionally, we observed a higher heart rate response in the stress group than in the control group (Figure 2B; Group × Time interaction; F(6,240) = 9.77; p<0.001). The Bonferroni post-hoc test revealed that these differences were significant at the anticipation (*t* = 3.014, p<0.05), speech (t = 5.736, p<0.001) and math (t = 7.00, p<0.001) time points (Figure 2b). The salivary cortisol concentration also increased significantly after the stress protocol in comparison with the control group (Figure 2C; Group × Time interaction; F(4,160) = 8.793; p<0.001); using the Bonferroni post-hoc test, we found that this increase was particularly significant at sampling time +10 (*t* = 2.909, p<0.05) and +25 (t = 2.740, p<0.05). In conclusion, these results showed that the stress (TSST) and control protocols were properly implemented.

**Figure 2.**
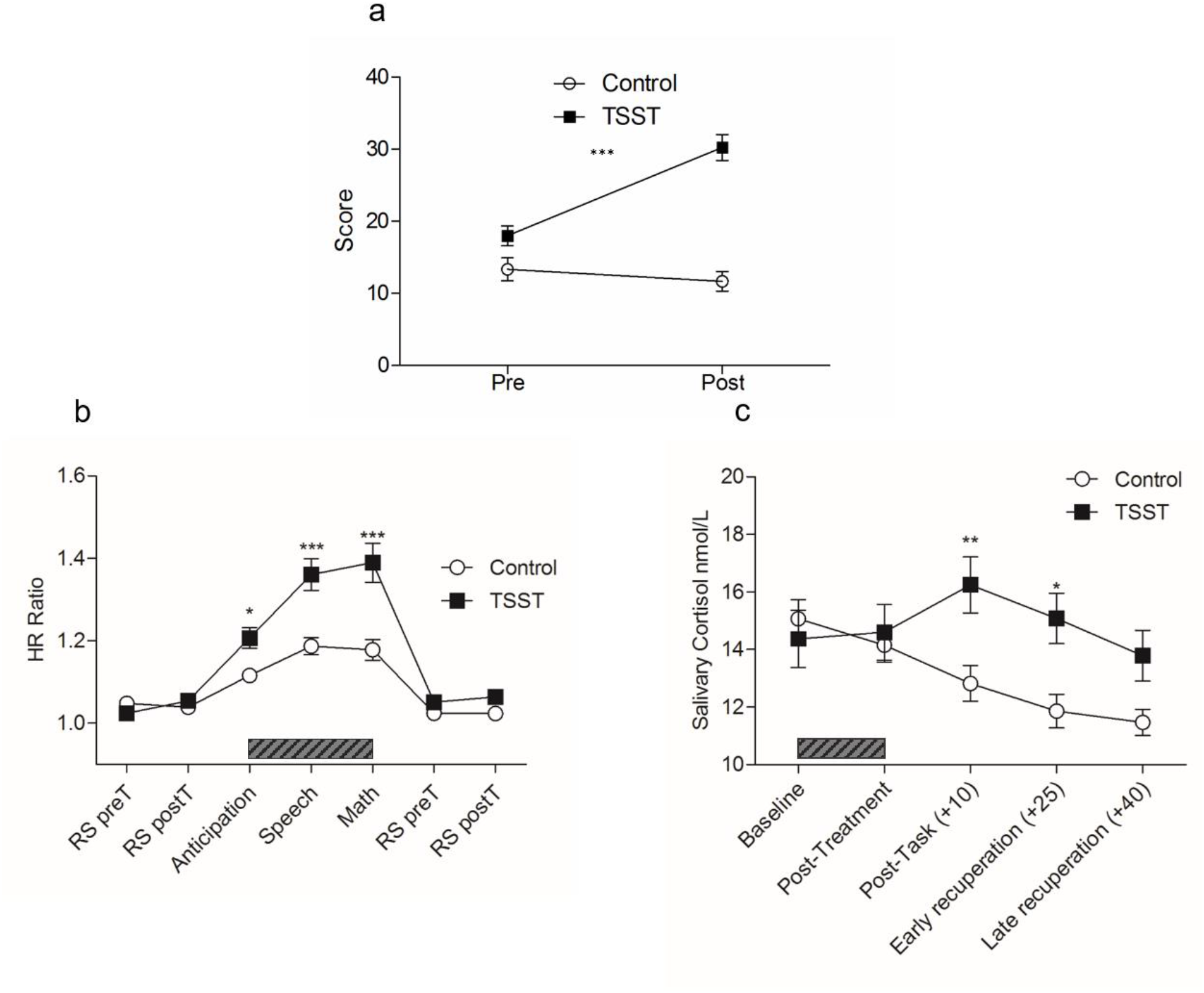
Psychological and physiological stress responses to the Trier Social Stress Test (TSST) or the control protocol. a) The self-reported level of anxiety was calculated by subtracting the baseline measurement from the score on the level of anxiety scale after the TSST or the control protocol. b) The mean heart rate during two 90-second rest periods flanking the baseline attentional task, during the 90-second anticipation, speech and math periods of the TSST or the control protocol, and during the 90 seconds of rest flanking the attentional task after treatment (RS pre-T: Resting state pre-task, RS post-T: Resting state post-task). c) The salivary cortisol concentration was measured at five points during the protocol: just after setup (baseline), just after treatment (Time = 0), and after 10, 25 and 40 minutes of the protocol. The gray bars in b and c indicate either the TSST or the control protocol. The error bars represent the standard errors of the means (SEM). There were N = 21 participants per group. *p<0.05; **p<0.01; ***p<0.001.

Finally, the participants reported no differences in their trait anxiety (Figure S1a, p>0.05) or in their daily levels of subjective perceived stress (Figure S1b, p>0.05). Therefore, we can assume that the results of this report are due to the experimental protocol rather than external daily stress or intrinsic personality features of the participants.

### Attentional performance improvement is affected by psychosocial stress

We defined accuracy as the number of correct trials and measured the reaction time before and after both protocols. Psychosocial stress impairs performance improvement, which was observed after the control protocol (Figure 3A; Group × Time interaction; F(1,40) = 5.870; p<0.05). However, a similar decrease in the reaction time for correct answers was observed after both protocols (Figure 3B; Time × Group interaction; F(1,40) = 0.002; p>0.05; Time main effects; F(1,40) = 35.87; p<0.001), which suggests that the effect of stress on performance was not due to fatigue or poor attitude toward the task. Moreover, it may be possible that when giving correct answers, the participants of both groups were equally attentive.

**Figure 3.**
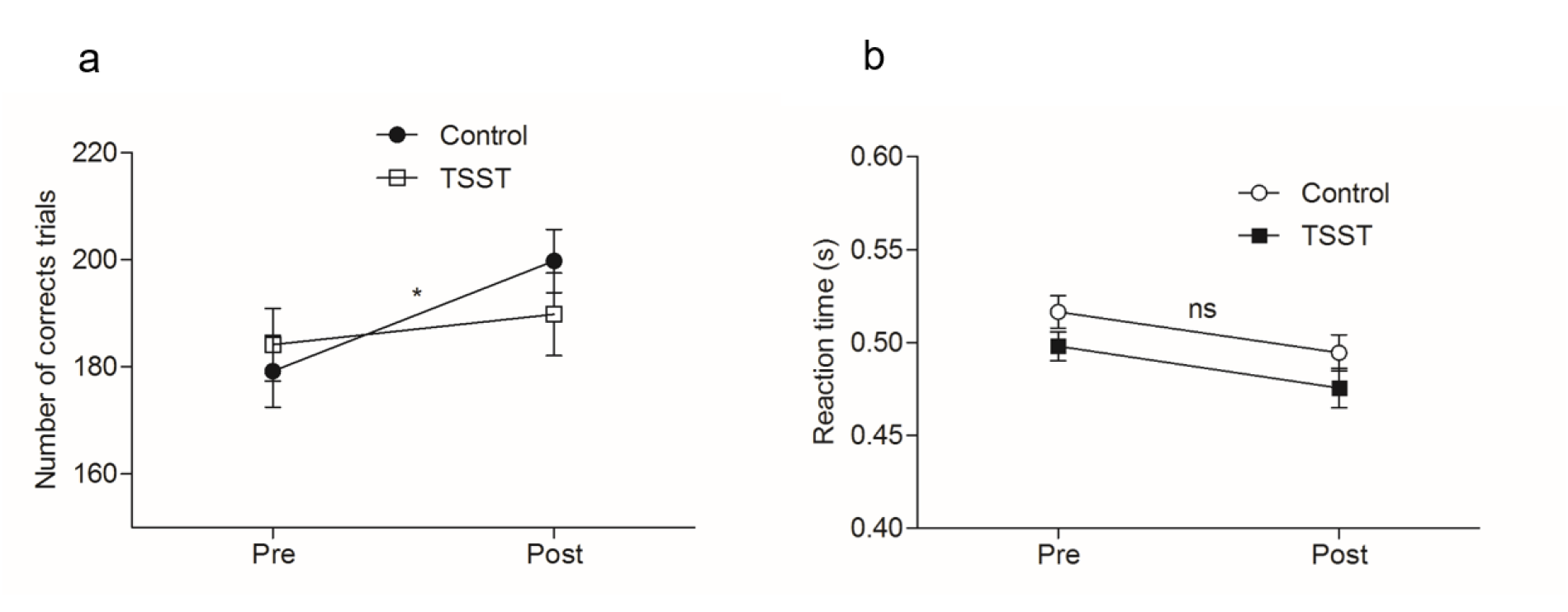
Effects of psychosocial stress on attentional performance. Attentional accuracy (the number of correct responses trials) (a) and reaction times (b) were measured during the attentional task before (pre) and after (post) treatment. The error bars represent the standard errors of the means (SEM). There were N = 21 participants per group; *p<0.05; ns, nonsignificant.

### Attentional performance varies with the level of anxiety acquisition

After performing a Pearson correlation analysis of the change in attentional performance and both physiological activation and the self-reported level of anxiety, we observed that the only stress outcome that correlated with attentional performance was anxiety; the increase in anxiety corresponded to the decrease in attentional performance (Table S1, Figure 3a). To explore the negative relationship between attentional performance and anxiety, we separated the TSST group into sub-groups with low and high levels of anxiety acquisition using the median as a cut point (Figure 3) and compared the attentional performance for the 3 groups (control, low anxiety TSST and high anxiety TSST). Interestingly, the participants with low levels of anxiety acquisition (mean, 5.9 ± 2.002; n = 10) behaved similarly to the participants in the control group (mean, −0.6 ± 0.89; n = 21) (Figure 3b; Group × Time interaction; F(1,29) = 0.03; p>0.05). In those with high levels of anxiety acquisition (mean, 17.91 ± 0.9091; n = 11), the attentional performance not only failed to improve but became worse; as it differed significantly from results for the participants with low anxiety acquisition and the participants in the control group (Figure 3b; Group × Time interaction; F(1,19) = 9.081; p<0.01). The same procedure was performed for participants with high and low cortisol levels and heart rates; however, there were no differences in performance between the low and the high responders in any of the cases (Figure S2a-b).

### Relationship between attentional performance and stress response

We used the MATLAB toolbox (M3) to study the relationships between different stress variables and attentional performance for the 42 participants. Table 1 shows the path coefficient, standard error and significance for each different single-level model. We found that anxiety was negatively correlated with attentional performance independent of changes in the cortisol level (Table 1A, c’, ab) and heart rate (Table 1B, c’, ab), nevertheless, changes in the cortisol level (Table 1C, a) and heart rate (Table 1E, a) were inversely related to changes in anxiety (Table 1A, B, a). Interestingly, neither the cortisol level (Table 1C, c’) nor the heart rate (Table 1E, c’) was related to attentional performance individually; however, there was a significant effect involving anxiety as a mediator for both (Table 1C, E, ab). Finally, the cortisol level and the heart rate were mutually related (Table 1D, F, a).

**Table 1.** Mediation analysis values

| Seed/Mediator | a | b | $c'$ | ab |
| --- | --- | --- | --- | --- |
| A) Anxiety/cortisol | 0.17** | 0.24 | -1.19** | 0.04 |
| B) Anxiety/HR | 0.02*** | 6.76 | -1.32** | 0.16 |
| C) Cortisol/anxiety | 0.77* | -1.19** | 0.24 | -0.93* |
| D) Cortisol/HR | 0.04* | -1.34 | -0.71 | 0.01 |
| E) HR/anxiety | 8.04** | -1.31** | 6.69 | -10.83** |
| F) HR/cortisol | 3.88* | -0.69 | -1.33 | -2.83 |

Each box includes the path coefficient for each single-level model considering the 42 participants.

### Discussion

We proposed an experiment designed to study the relationship between three different landmarks of psychosocial stress, cortisol level, heart rate and anxiety, and performance in an attentional task. Our hypotheses were confirmed and are summarized as follows: (1) acute psychosocial stress resulted in failure to improve performance in the attentional task, which was clearly observed after the control protocol (Figure 3); (2) stress-related physiological responses, including increases in both salivary cortisol level and heart rate, were positively correlated with the self-reported level of anxiety (Table S1); (3) the effect of psychosocial stress on attentional control was directly related to the self-reported level of anxiety augmentation (Table 1). In particular, the participants in the stress group with high anxiety acquisition performed less well after the TSST compared to their baseline performance, whereas those with low anxiety acquisition improved their performance as much as the participants in the control group did (Figure 4). Therefore, our results highlight the relevance of the stress-dependent self-reported level of anxiety to attentional control.

**Figure 4.**
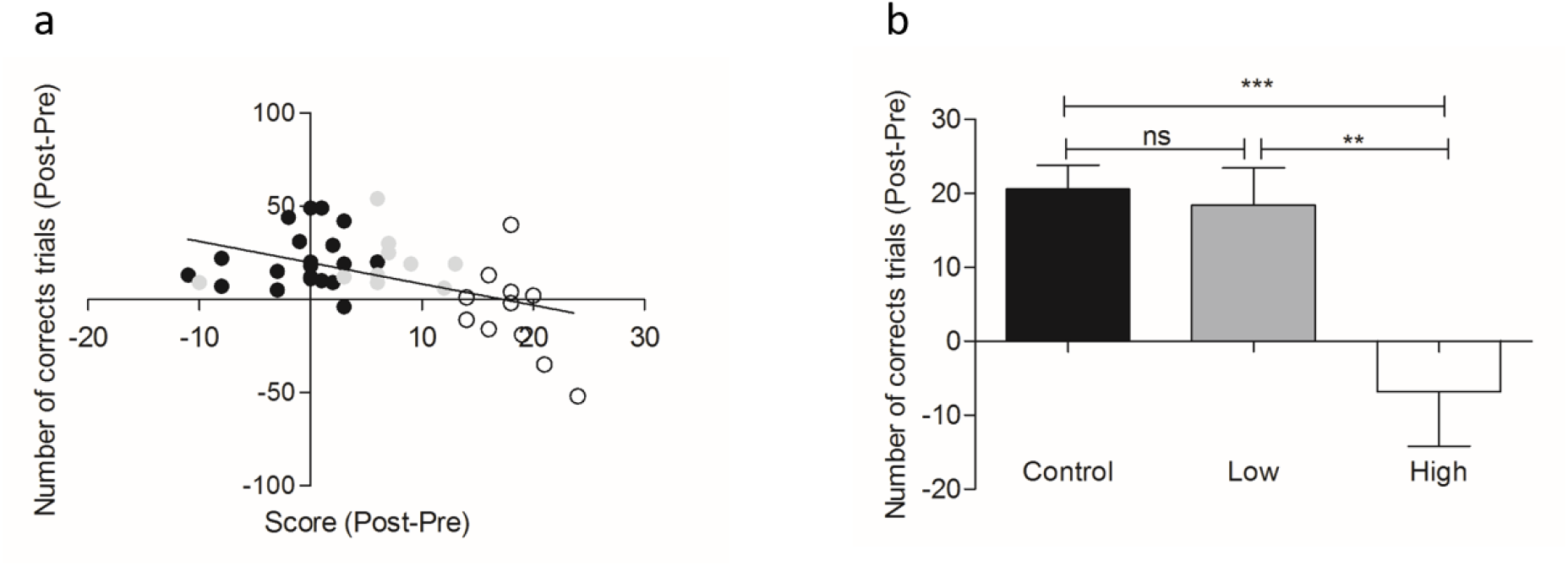
Dependence between attentional performance and level of anxiety. a) Scatter plot of the entire sample divided among the control (black circles), low anxiety acquisition (gray circles) and high anxiety acquisition (white circles) groups. b) Attentional accuracy was measured by the number of correct responses after treatment and was corrected using the baseline of the control and low and high anxiety groups (n: control=21, low=10, high=11). The error bars represent the standard errors of the mean (SEM). **p<0.01; ***p<0.001.

According to classical mediation (Baron & Kenny, 1986), none of the six single-level models tested satisfied the four requirements for establishing mediation (Methods/ Single-level model). The single-level model that came closest to satisfying those requirements was tested using the level of anxiety as the mediator variable, the salivary cortisol level or the heart rate as the initial variable (Table 1, C, E) and attentional performance as the outcome variable. Both single models failed on the first requirements because neither the cortisol level nor the heart rate was statistically related to the attentional performance even though contemporary approaches have proposed that the first requirement is not mandatory as long as the indirect effect carried by the relationship between X and M and M and Y was present (MacKinnon, Lockwood, Hoffman, West, & Sheets, 2002). Nevertheless, this type of relationship, known as indirect effect, is not considered classical mediation (Mathieu & Taylor, 2006). Accordingly, our analyses suggest that both salivary cortisol level and heart rate are indirectly related to attentional performance through their relationship with the self-reported level of anxiety. There is an indirect effect between physiological response and attentional performance through the self-reported level of anxiety. Interestingly, in our single-level models (Table 1, C, E), the path coefficients *c*’ and *ab* had opposite signs. This phenomenon, called inconsistent mediation, occurs when the supposed mediator variable M acts as a suppressor variable (MacKinnon, Fairchild, & Fritz, 2007).^2^

The observed relationship between the self-reported level of anxiety and attentional performance is consistent with the findings of Abercrombie, Speck, and Monticelli (2006), who highlighted the relevance of the psychological perception of stress to memory. This hypothesis is also in accordance with the study of Liston et al. (2009), who showed that as long as daily stress perception increased, attentional performance did not. However, other studies have found no interactions among stress level, psychological state and task performance (Plessow et al., 2012; Schoofs et al., 2008).

Analogously, we did not find any direct relationship between attentional performance and cortisol response. This could be because the second attentional task was performed immediately after the stress protocol when the salivary cortisol level had not yet peaked (Figure 2C). The time lag between a stressor and cortisol activity is approximately 30 minutes (Sapolsky, Romero, & Munck, 2000). This issue was discussed by Het, Ramlow, and Wolf (2005) in the field of memory, in which, apparently, the timing of the treatment with respect to the course of the study (before learning vs. before retrieval) is important for evaluating the effect of stress on memory. Similarly, studies in which performance was evaluated when the cortisol concentration was elevated showed a negative correlation between behavioral performance and the cortisol response (Clemens Kirschbaum, Wolf, May, Wippich, & Hellhammer, 1996; Roelofs et al., 2007; Schoofs et al., 2008). Furthermore, the effect of psychosocial stress on cognitive function was not present immediately after stress induction but gradually became evident (Plessow, Fischer, Kirschbaum, & Goschke, 2011). Consistent with our results, some researchers have found no correlation between cortisol levels and performance even in tasks performed at peak cortisol concentration (Cornelisse, van Stegeren, & Joels, 2011). The lack of relationship may be explained as follows: at the time of the second attentional task (Figure 1), the increase in the cortisol level was not accompanied by an increase in the autonomic response measured by an increase in heart rate (Figure 2b and c). This has been described as necessary for working memory impairments (Elzinga & Roelofs, 2005). Apparently, the autonomic stress response is as involved in stress-dependent alterations as the cortisol response (Chamberlain et al., 2006).

The interaction between physiological and psychological responses to psychosocial stress is definitely an issue that has not been understood until now. Our analysis showed a positive reciprocal interaction between cortisol reactivity, heart rate increase and anxiety acquisition. The results are consistent with those of Schlotz, Schulz, Hellhammer, Stone, and Hellhammer (2006), in which the relationship between cortisol and performance is explained by changes in trait anxiety, yielding a stronger relationship in the subjects with higher trait anxiety. In contrast, high trait anxiety was associated with lower neuroendocrine reactivity during psychosocial stress (Jezova, Makatsori, Duncko, Moncek, & Jakubek, 2004), with the result that higher anxiety was associated with an inability to respond to stress. A recent study, which aimed to explore the relationship between physiological and psychological stress responses, concluded that the emotional experience of stress and the physiological stress response were two dissociated systems (Ali et al., 2017). This result contrasts with what we found; it is likely that the level of anxiety was a more reliable psychological stress marker than the one used in the mentioned work, i.e., mood. Interestingly, the same authors discussed the likelihood that the commonly used psychological stress markers could be not valid and appropriate measures, especially at discrete times, for studies of their relationships with body responses in the context of psychosocial stress.

Theoretically, the dynamic model of stress and sustained attention of Hancock and Warm (2003) proposes that stress decreases the available attentional capacity, which can be explained as psychological adaptability. In this way, there could be a direct relationship among stress, psychological state and attention, which is consistent with our results and the results of others (Liston et al., 2009; Vedhara, Hyde, Gilchrist, Tytherleigh, & Plummer, 2000). Interestingly, Eysenck et al. (2007) proposed that elevated states of anxiety affect attentional control, disrupting the top-down/bottom-up balance; therefore, attention is mainly allocated to threat-related stimuli. Following this idea, it was shown that anxiety was associated with increased attention to threatening images (Quigley, Nelson, Carriere, Smilek, & Purdon, 2012; van Honk et al., 2001). Furthermore, another study showed that a very anxious participant can reorder images of angry faces (compared to neutral) that have presented and then, placed in different locations more effective, which provides evidence of increased emotional memory due to anxiety (Terburg, Aarts, & van Honk, 2012). Our attentional task lacks an emotional or threating stimulus; however, if we consider the psychosocial stress experience a threat or challenge, then, we expect increased attention to this experience even when it occurred in the past. Because attentional capacity is limited, the allocation of attentional resources to subjects other than the task (external task-irrelevant distractors or internal thoughts relating to previous stressful experiences) may diminish the likelihood of performing the task well.

Our study was restricted to men because it has been shown that the menstrual cycle and the use of oral contraceptives can strongly affect the stress neuroendocrine response in women (C. Kirschbaum, Kudielka, Gaab, Schommer, & Hellhammer, 1999). Moreover, it has also been shown that the relationship between the subjective stress response and the cortisol level depends on the phase of the menstrual cycle (Duchesne & Pruessner, 2013). Therefore, the results of this study cannot be extended to women. More studies with larger and more extensive samples are required.

Finally, some researchers have begun to explore the neural correlates associated with stress, anxiety and attentional control. One study showed a negative relationship between oscillatory brain activity at delta (4–6 Hz) and beta (13–29 Hz) frequencies and anxiety-driven attentional avoidance (Putman, 2011). Moreover, because beta-frequency activity is related to goal-directed top-down processes (Bastos et al., 2015; Buschman & Miller, 2007), we expect increased beta activity in situations in which increased top-down control is required (e.g., after psychosocial stress). Additional studies that explore the relationships among beta activity, anxiety and attentional control will enhance our understanding of how social stress affects behavior.

### Conclusion

The results of the present study highlight the relevance of immediate stress-dependent anxiety acquisition to understanding the effects of social stressful situations on attentional control. The subjective experience of anxiety is apparently more relevant than the physiological stress response, which could play an indirect role by affecting the perception of the current body state. This finding is relevant because it returns the focus to the subjective psychological experience rather than stress-induced involuntary physiological changes. Such an observation may have important implications for designing therapeutic interventions to address social stress and stress in general. It could be far better to develop focused strategies to address psychological self-perception in stressful environments than to attempt to reduce the natural stress-induced physiological reaction, which could be used to benefit the subject.

## Acknowledgments

This work was supported by a doctoral fellowship (CONICYT 21140884, Chile) awarded to I.P., grants from FONDECYT awarded to J. R. S. (1130810, Chile) and E. R. (1120752, Chile), and a grant from the Fund for Innovation and Competitiveness of the Chilean Ministry of Economy, Development and Tourism through the Millennium Scientific Initiative (IS130005) awarded to J. R. S. and E. R.

## Footnotes

1 Classical mediation has 4 requirements: (1) an initial relationship between X and Y, (2) a relationship between X and M, (3) a relationship between M and Y, and (4) a decrease in the relationship between X and Y in the presence of M.

2 Both cortisol and heart rate may have positive relationships with attentional performance (Table 1 C, E, c); however, in the presence of anxiety, these relationships become negative (Table 1 C, E, c’ + ab). Therefore, an acute increase in the physiological response may have beneficial effects on attention as long as the level of anxiety does not increase. Additional research on this relationship are necessary to unravel this issue.

